# Co-outbreak of ST37 and a novel ST3006 *Klebsiella pneumoniae* from multi-site infection in a neonatal intensive care unit: a retrospective study

**DOI:** 10.1101/334169

**Authors:** Dongjie Chen, Xinlan Hu, Falin Chen, Hongru Li, Daxuan Wang, Xiaoqin Li, Changsheng Wu, Ning Li, Shaolian Wu, Zhen Li, Liqing Chen, Yusheng Chen

**Affiliations:** Shengli Clinical Medical College of Fujian Medical University, Fuzhou, 350001, China; Clinical Microbiology Laboratory, Fujian Provincial Hospital, Fuzhou 350001, China; Department of Respiratory Medicine and Critical Care Medicine, Fujian Provincial Hospital, Fuzhou 350001, China

**Author notes:** Corresponding author: Yusheng Chen, No.134 East Street, Fuzhou 350001, China.

**Keywords:** Co-outbreak, *Klebsiella pneumoniae*, Multidrug resistance, Genome sequencing, Neonatal intensive care unit

## Abstract

**Background:** The outbreak of carbapenems resistant *Klebsiella pneumoniae* (K. pneumoniae) is a serious public health problem, especially in the neonatal intensive care unit (NICU).

**Methods:** Fifteen strains of *K. pneumoniae* were isolated from seven neonates during June 3–28, 2017 in a NICU. Antimicrobial susceptibility was determined by the Vitek 2 system and micro-broth dilution method. Multi-locus sequence typing (MLST) and pulsed-field gel electrophoresis (PFGE) were used to analyse the genetic relatedness of isolates. Genome sequencing and gene function analyses were performed for investigating pathogenicity and drug resistance and screening genomic islands.

**Findings:** Two *K. pneumoniae* clones were identified from seven neonates, one ST37 strain and another new sequence type ST3006. The ST37 strain exhibited multi-drug resistance genes and resistance to carbapenem. MLST and PFGE showed that 15 strains were divided into three groups, with a high level of homology. Gene sequencing and analysis indicated that KPN1343 harboured 12 resistance genes, 15 genomic islands and 205 reduced virulence genes. KPN1344 harboured four resistance genes, 19 genomic islands and 209 reduced virulence genes.

**Conclusion:** Co-outbreak of *K. pneumoniae* involved two clones, ST36 and ST3006, causing multi-site infection. Genome sequencing and analysis is an effective method for studying bacterial resistance genes and their functions.

## Introduction

*Klebsiella pneumonia*, particularly multidrug resistant *K. pneumoniae*, is a common and important pathogen in hospitals, posing a serious and urgent threat to public health (1, 2). Neonates in the neonatal intensive care units often have underlying conditions, such as prematurity, or the presence of indwelling catheters, a history of antibiotic treatment and parenteral nutrition, which are risk factors for infection (3). In addition, relaxed vigilance by doctors and nurses regarding nosocomial infection promotes nosocomial infection outbreaks, with considerable impact on neonatal treatment and prognosis, prolonged hospital stays, increased hospital costs and mortality.

Nosocomial outbreaks in the neonatal intensive care unit are frequently reported (4-7), however, it is rare to isolate the same clonal pathogen from different sites in the same patient. This study identified 15 *K. pneumoniae* strains isolated from sputum specimens, blood specimens and umbilical vein catheter tips in seven neonates that belonged to the two cloned strains ST37 and ST3006, which produced carbapenemase and extended-spectrum beta-lactamases, respectively.

## Methods

### Patients and bacterial strains

Patient characteristics were obtained from electronic medical records. Bacterial strains were isolated from seven neonates and stored in a refrigerator (SANYO, Japan) at −70°C. Strains were identified using the Vitek 2 system (BioMérieux, France). This study was approved by the ethics committee of hospital, the ethics approval number is K2018-01-001.

### Multi-locus sequence typing

Bacterial DNA was extracted using the Bacteria Genomic DNA Kit (CWBIO, China). Multi-locus sequence typing (MLST) for *K. pneumoniae* was performed following the methods described previously (8). The allelic profiles and sequence types (STs) were Determined using online databases (https://pubmlst.org/bigsdb?db=pubmlstmlstseqdef). The novel allele profiles were sent to for confirmation.

### Pulsed-field gel electrophoresis

Pulsed-field gel electrophoresis (PFGE) of Xbal-digested genomic DNA samples of *K. pneumoniae* was performed with a CHEF MAPPER XA apparatus (Bio-Rad, USA) according to the literature (9). Electrophoresis was performed for 24 h at 14°C with pulse time ranging from 5 to 35 s at 6 V/cm. PFGE profiles were analysed and compared using the Gel Doc XR+ system, version 2.0 (Bio-Rad, USA).

### Antimicrobial susceptibility testing

Antibiotic susceptibility testing was performed by the Vitek 2 system (BioMérieux, France), including ampicillin sulbactam, cefazolin, ceftriaxone, cefotetan, ceftazidime, cefepime, gentamicin, tobramycin, amikacin, levofloxacin, ciprofloxacin, aztreonam, imipenem, ertapenem, piperacillin-tazobactam, trimethoprim-sulfamethoxazole. The MICs of tigecycline and polymyxin B was determined by the micro-broth dilution method. All antibiotics were interpreted according to the Clinical and Laboratory Standards Institute (CLSI) document M100-S27 (10), except tigecycline and polymyxin B. For tigecycline and polymyxin B, the European Committee on Antimicrobial Susceptibility Testing (EUCAST) breakpoint (11) was used. *E. coli* ATCC 25922 and *P. aeruginosa* ATCC27853 were used as quality control.

### Genome sequencing and gene analysis

Whole-genome sequencing was performed on the Illumina HiSeq PE150 platform (Novogene Bioinformatics Technology Co., Ltd., Beijing, China). The Island Path-DIOMB program was used to predict the genomic islands. For these pathogenic bacteria, we used the Pathogen-Host Interactions Database (PHI) and the Comprehensive Antibiotic Resistance Database (CARD) to perform the pathogenicity and drug resistance analyses.

## Results

### Clinical characteristics of patients

Patient clinical characteristics were determined from the retrospective study of the electronic medical records and are presented in Table I. Subjects included five premature infants and two infants with hyperbilirubinaemia. All of the five premature infants were also diagnosed as low birth weight infants (two were very low birth weight infants) and with neonatal respiratory distress syndrome (NRDS). Three infants were delivered by caesarean section, two infants were delivered by midwives, one infant was delivered by forceps and only one infant was a normal delivery. The average age of the five premature infants was approximately 30 weeks, with an average body weight of 1.37 kg. Patient 1 was treated with piperacillin/tazobactam and meropenem, while the remaining patients were treated with piperacillin/tazobactam. Patient 1, Patient 2, Patient 3, Patient 4, Patient 6 and Patient 7, had a lower than normal Apgar score (10-10-10). The clinical condition of Patient 4 (the second of the twins) was poor but was discharged upon request of the family, while the remaining patients were treated and discharged.

**Table I.** Clinical characteristics of patients

| Patients | Clinical diagnosis | Mode of delivery | Age (w) | Sex | Body weight (kg) | Antibiotics | Apgar score | Outcome |
| --- | --- | --- | --- | --- | --- | --- | --- | --- |
| P1 | Premature infant<br>Low birth weight infant<br>NRDS | Caesarean delivery | 33 | F | 1.55 | TZP,<br>MEM | 6-8-8 | Treated |
| P2 | Premature infant<br>Low birth weight infant<br>NRDS | Normal delivery | 31 | F | 1.60 | TZP | 8-8-8 | Treated |
| P3 (first of twins) | Premature infant<br>Very low birth weight infant<br>NRDS | Midwifery delivery | 29 | F | 1.45 | TZP | 8-9-9 | Treated |
| P4 (second of twins) | Premature infant<br>Very low birth weight infant<br>NRDS | Midwifery delivery | 29 | F | 1.26 | TZP | 8-9-9 | Automatic discharge |
| P5 | Neonatal hyperbilirubinemia | Forceps delivery | 40 | M | 2.76 | TZP | 10-10-10 | Treated |
| P6 | Neonatal hyperbilirubinemia | Caesarean delivery | 40 | F | 3.83 | TZP | 10-10-10 | Treated |
| P7 | Premature infant<br>Very low birth weight infant<br>NRDS | Caesarean delivery | 30 | M | 1.00 | TZP | 6-8-8 | Treated |

P: patient; NRDS: neonatal respiratory distress syndrome; W: weeks; F: female; M: male; kg: kilogram; TZP: piperacillin/tazobactam. MEM: meropenem.

### Characteristics and Antibiotic Susceptibility of *K. pneumoniae*

Twenty-one *K. pneumoniae* strains were identified by the VIETK 2 system from different specimens, including sputum, blood, umbilical vein catheter tip. Only 15 *K. pneumoniae* strains were collected and frozen in a −70°C refrigerator. In Patient 2, Patient 3, Patient 4 and Patient 6, we simultaneously isolated two *K. pneumoniae strains* from blood and/or umbilical vein catheter tips and ST typing and drug resistance were consistent. (ST37, a novel ST typing ST3006: gapA 69, infB 19, mdh 90, pgi 20, phoE 125, rpoB 18, novel allele tonB 406). Antibiotic susceptibility testing showed that the ST37 strain was resistant to ampicillin-sulbactam, cefazolin, ceftriaxone, cefotetan, ceftazidime, cefepime, gentamicin, aztreonam, imipenem, ertapenem, piperacillin-tazobactam. The ST3006 strain was only resistant to ampicillin-sulbactam, cefazolin, ceftriaxone. (Table II, Table III). After investigation and nosocomial infection control are carried out by the infection department, no further bacteria were isolated from the patients, except for a strain ST1224 from Patient 3 on June 28, 2017.

**Table II.**
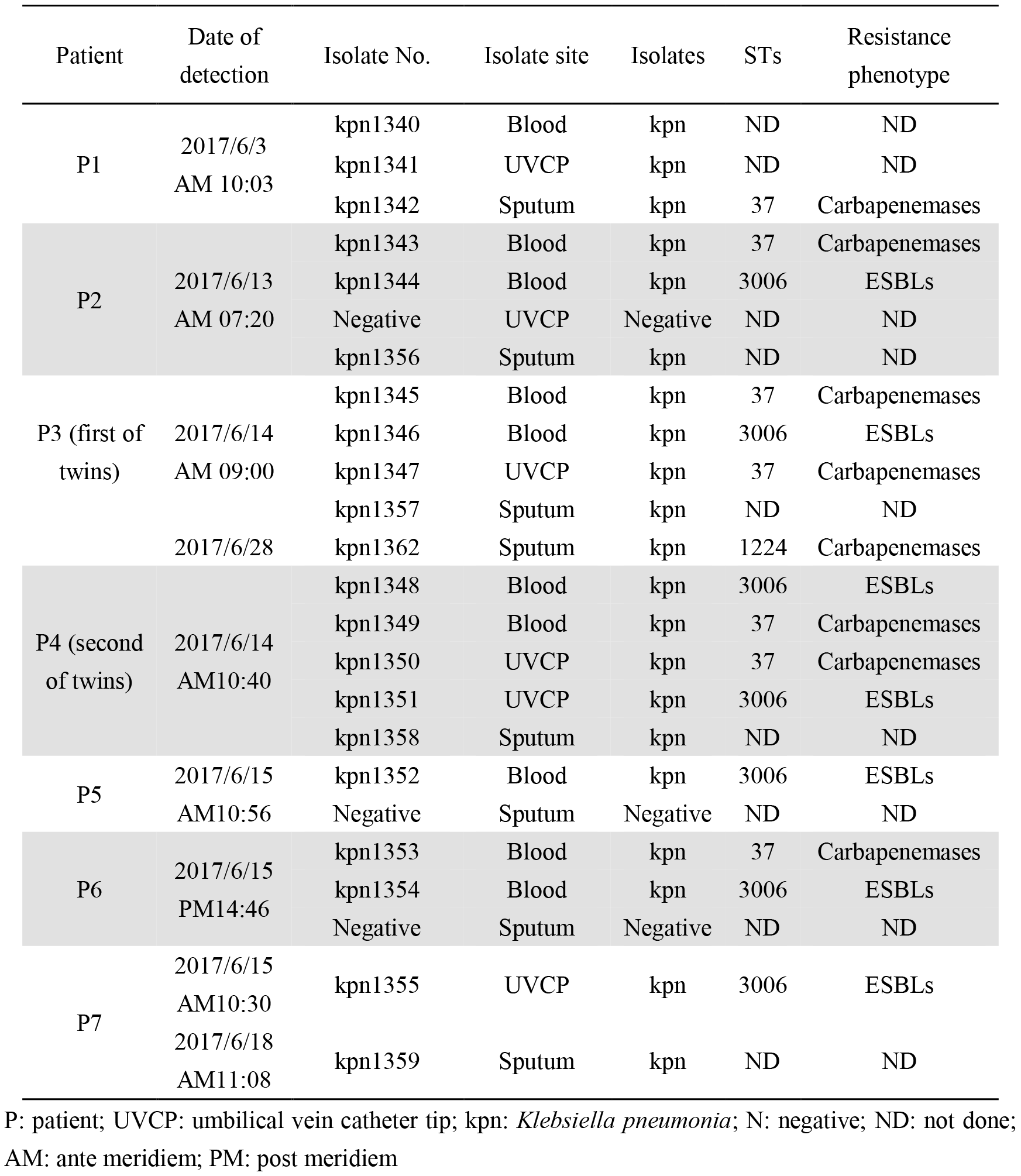
The characteristics of *K. pneumoniae*

**Table III.** Antibiotic susceptibility of *K. pneumoniae* Isolates ST37 and ST3006

| Antibiotics | MIC (ug/mL) |  | S/I/R |  |
| --- | --- | --- | --- | --- |
|  | *K. pneumoniae* |  | *K. pneumoniae* |  |
|  | ST37(n=7) |  | ST3006(n=7) |  |
| Ampicillin-sulbactam | ≥32/ | R | ≥32 | R |
| Cefazolin | ≥64/ | R | ≥64 | R |
| Ceftriaxone | ≥64 | R | ≥64 | R |
| Cefotetan | ≥64 | R | ≤4 | S |
| Ceftazidime | ≥64 | R | ≤1 | S |
| Cefepime | ≥64 | R | ≤1 | S |
| Gentamicin | ≥16 | R | ≤1 | S |
| Tobramycin | =8 | I | ≤1 | S |
| Amikacin | ≤2 | S | ≤2 | S |
| Levofloxacin | ≤0.25 | S | ≤0.25 | S |
| Ciprofloxacin | ≤0.25 | S | ≤0.25 | S |
| Aztreonam | ≥64 | R | =64 | R |
| Imipenem | =4 | R | ≤1 | S |
| Ertapenem | ≥8 | R | ≤0.5 | S |
| Tigecycline* | ≤0.25 | S | ≤0.25 | S |
| Polymyxin B* | ≤0.25 | S | ≤0.25 | S |
| Piperacillin–Tazobactam | =64 | R | ≤4 | S |
| Trimethoprim–Sulfamethoxazole | ≤20 | S | ≤20 | S |

MIC: minimal inhibitory concentration; S: sensitive; I: intermediary; R: resistance; \*: tigecycline and polymyxin B antibiotic susceptibility were determined by micro-broth dilution method

### Antibiotic Resistance Genes

The CARD was used to search for the names of resistance-related genes. KPN1343 was found to have 12 antibiotic resistant genes: beta-lactam resistance (OXA-33, TEM-1, SHV-11), aminoglycoside resistance [AAC (6’)-IId, AAC (3)-IIa, AAC (6’)-Ib-cr], phenicol resistance (catB3), rifamycin resistance (arr-3), sulfonamide resistance (sul1), efflux pump complex or subunit conferring antibiotic resistance (oqxB, oqxA, CRP), streptogramin resistance (catB3). KPN1344 was found to have four antibiotic resistant genes: beta-lactam resistance (TEM-1, CTX-M-3), efflux pump complex or subunit conferring antibiotic resistance (vgaC, CRP). We chose the resistance gene with a best-identities rate ≥0.99 (Table IV).

**Table IV.**
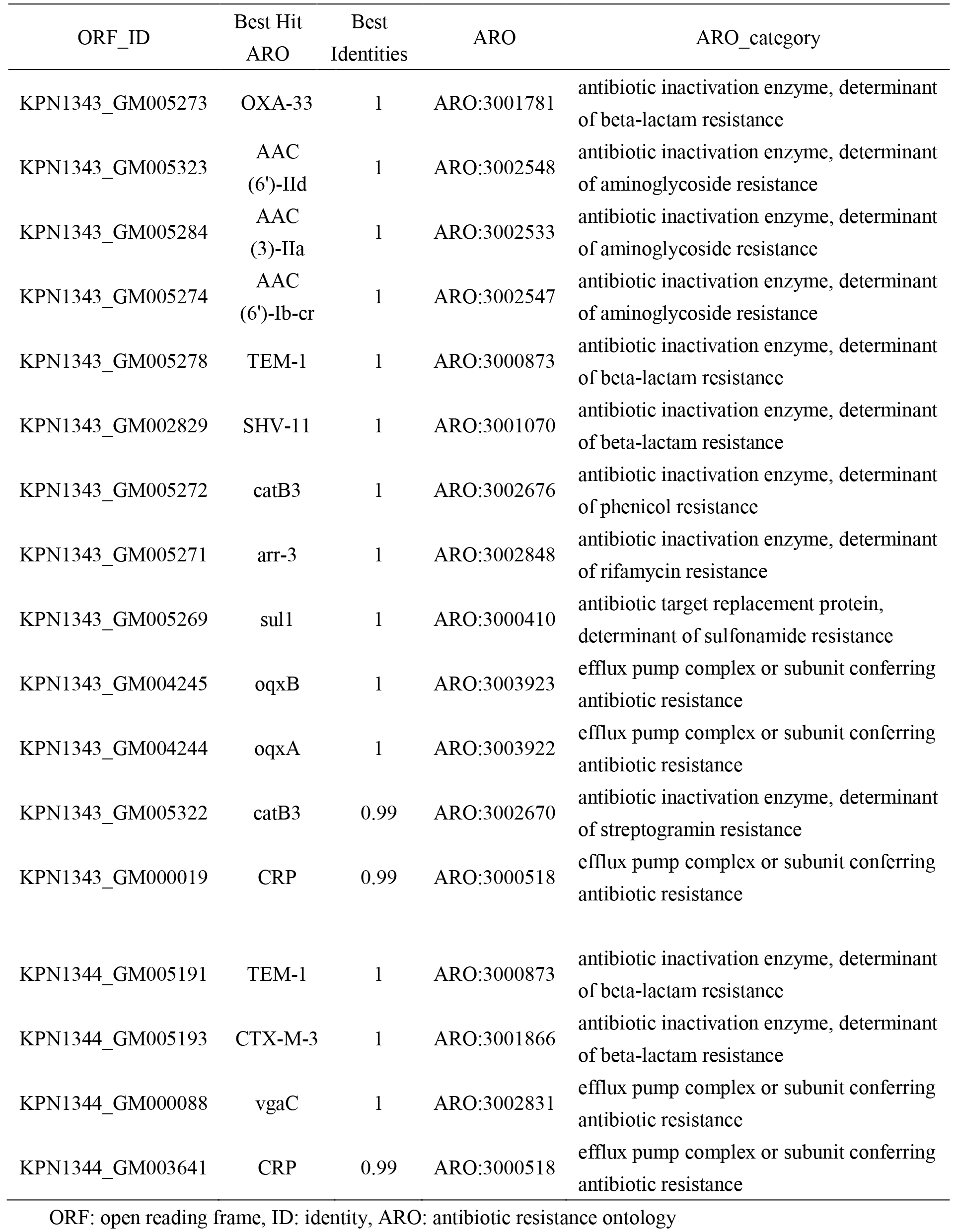
Antibiotic resistance ontology annotates for antibiotic resistance genes

### PFGE

PFGE homology analysis showed that 15 *K. pneumoniae* strains were divided into three clusters: ClusterA:1344,1346,1348,1351,1352,1354,1355; ClusterB:1362 and ClusterC:1342,1343,1345,1347,1349,1350, 1353. MLST typing also divided 15 *K. pneumoniae* strains into three groups. (Fig. 1)

**Fig. 1.**
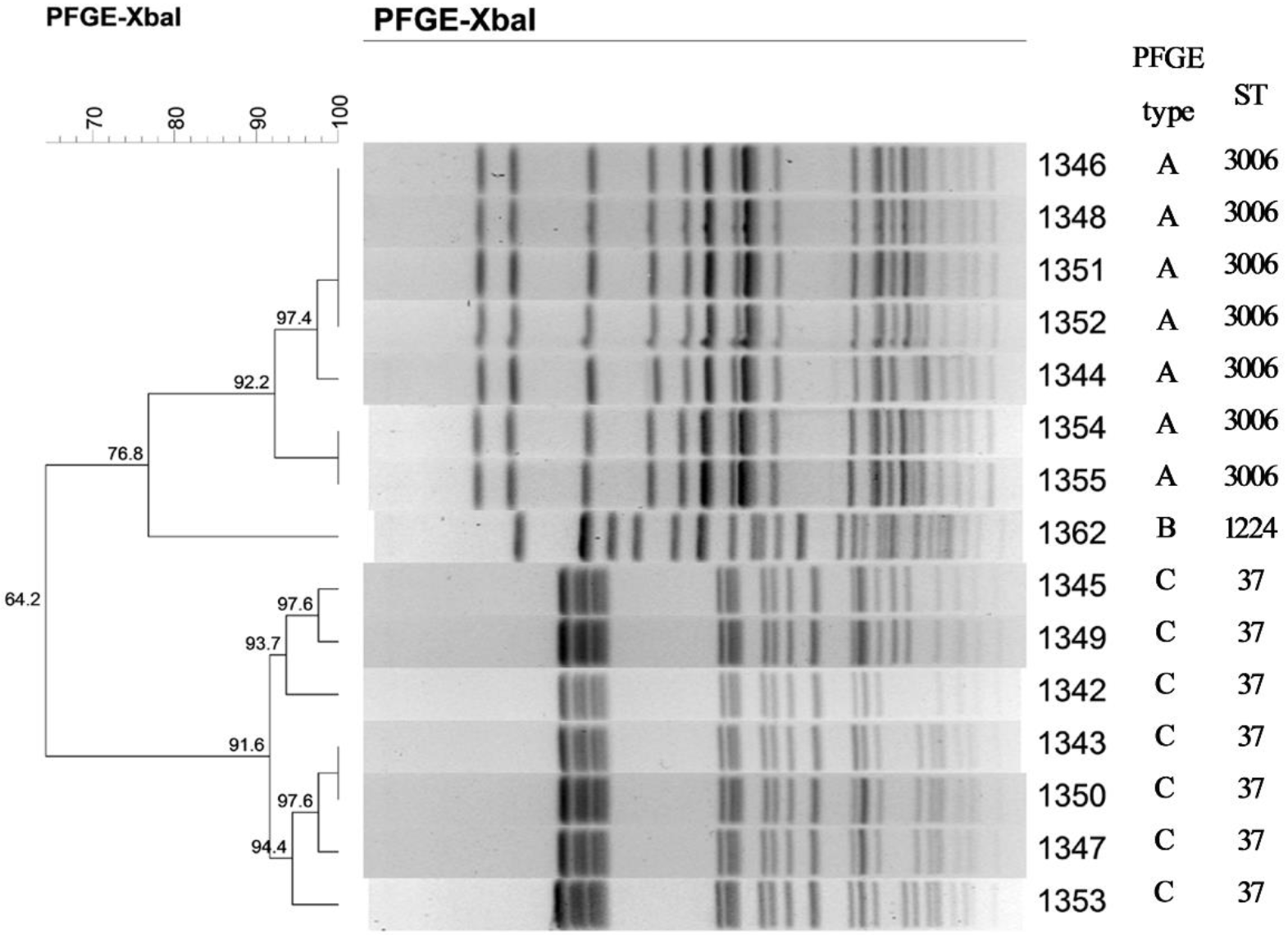
Pulsed-field gel electrophoresis (PFGE) of *K. pneumoniae*. 15 *K. pneumoniae* strains were divided into three clusters: Clusters A: 1344, 1346, 1348, 1351, 1352, 1354, 1355; Cluster B: 1362; Cluster C: 1342, 1343, 1345, 1347, 1349,1350, 1353. MLST typing also divided 15 *K. pneumoniae* strains into three groups: ST37, ST1224, ST3006. MLST: multi-locus sequence typing; ST: sequence type.

### Pathogen-Host Interactions and Genomic Islands

Using the BLAST software, the amino acid sequences of KPN1343 and KPN1344 were compared using the PHI database. PHI Phenotype classification showed that the number of reduced virulence genes matched the database mostly, 205 and 209, respectively (Fig. 2). Using Island Path-DIOMB to predict genomic islands, KPN1343 has 15 genomic islands and KPN1344 has 19 genomic islands, the length and direction of the genes are shown in Fig. 3 (length of <15 kb only is shown).

**Fig. 2.**
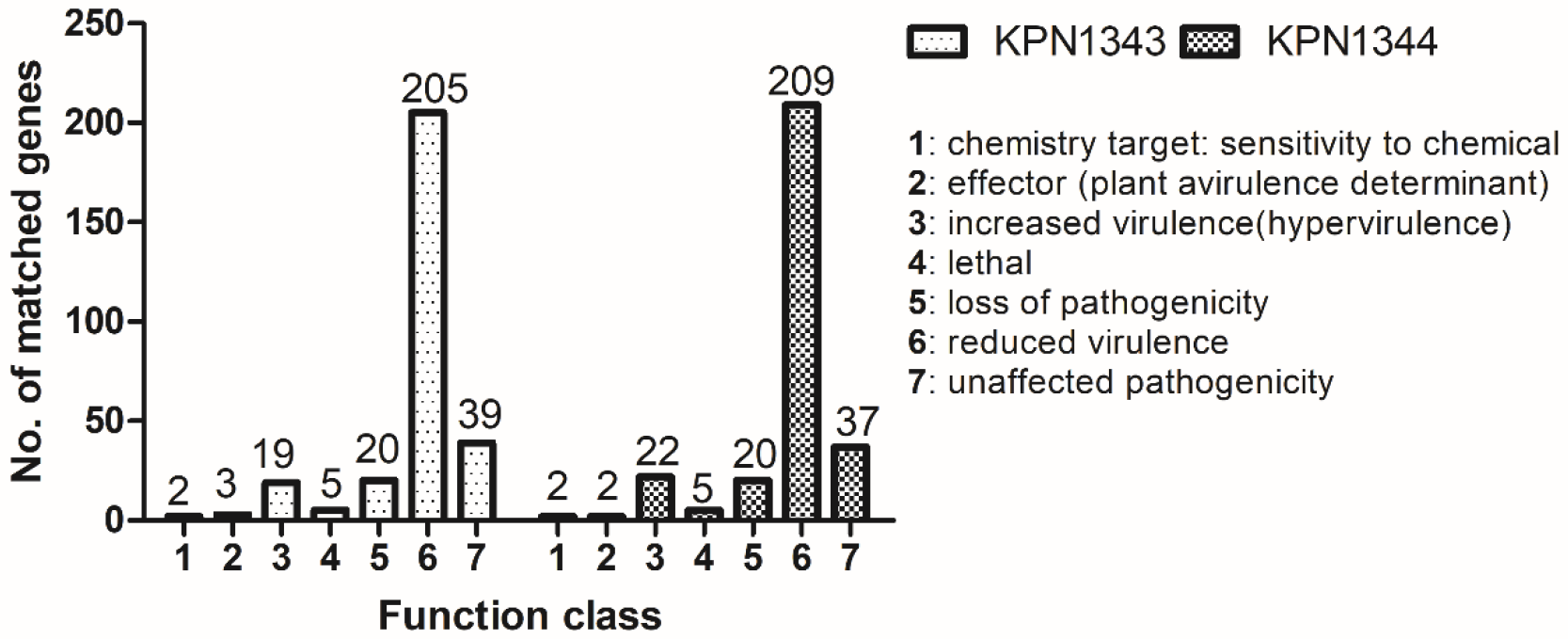
KPN1343 and KPN1344 PHI Phenotype classification. PHI Phenotype classification showed that the number of reduced virulence genes matched the database mostly, 205 and 209, respectively. PHI: pathogen-host interactions database.

**Fig. 3.**
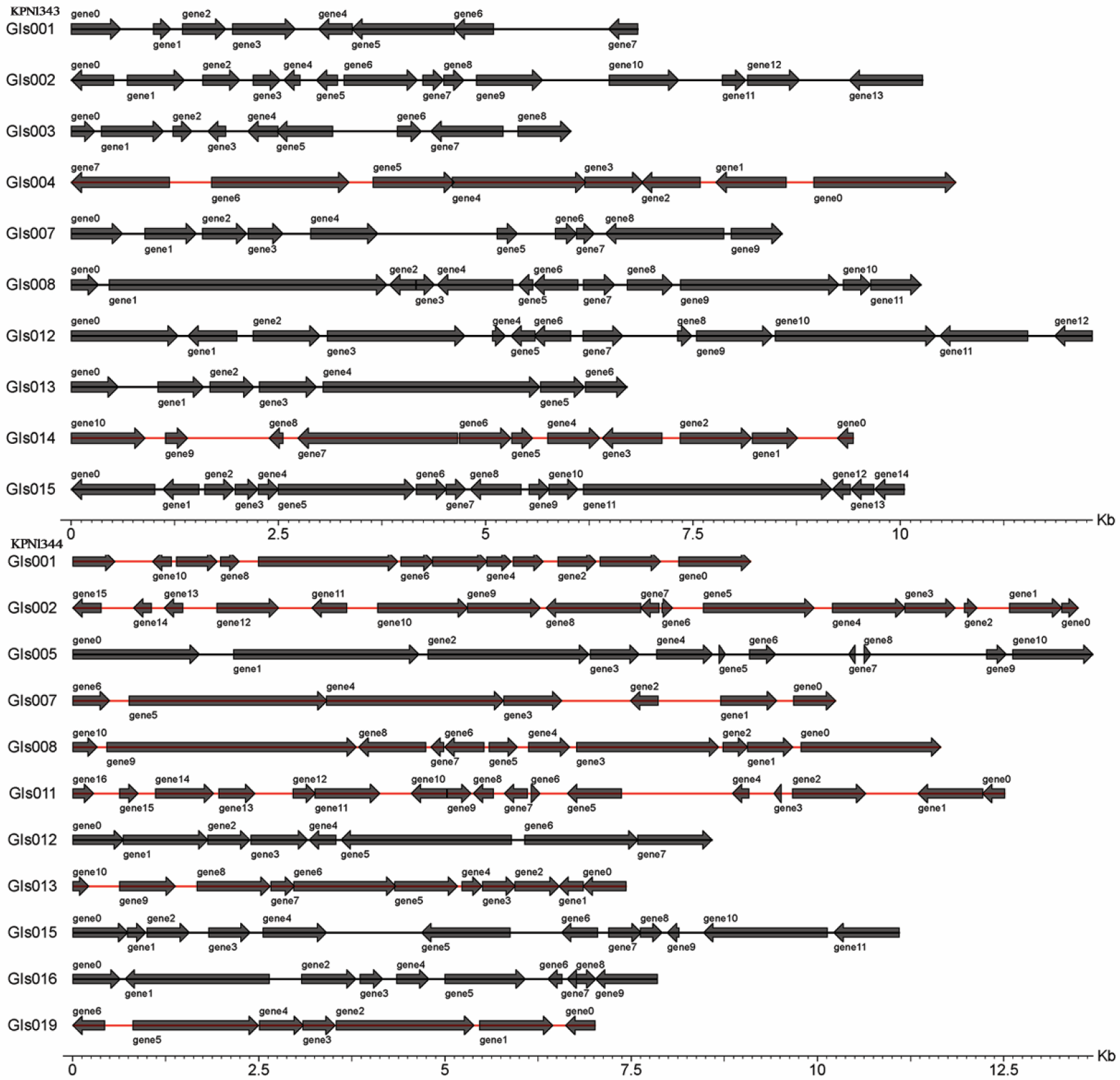
KPN1343 and KPN1344 genomic islands. KPN1343 has 15 genomic islands and KPN1344 has 19 genomic islands, the length and direction of the genes are shown in Fig. 3 (length of <15 kb only is shown).

## Discussion

*K. pneumoniae* is an important nosocomial pathogen that can cause pneumonia, urinary tract infection, digestive tract infection, bloodstream infection, liver abscess and meningitis (12-17). Infection outbreaks have been frequently reported in neonatal intensive care units. Extended spectrum beta-lactamase-producing and carbapenemase-producing *K. pneumoniae* can cause large outbreaks with significant morbidity and mortality effects (18). Underlying conditions, including premature delivery, low birth weight and neonatal respiratory distress syndrome, are risk factors for neonatal infection. Inadequate medical equipment, environmental disinfection and hand hygiene as well as understaffing have been reported to be important factors for infection outbreak (19, 20). In our study, infection control practitioners took samples from incubators, air, the hands of doctors and nurses, objects and patient skin for bacterial culture. Only one *K. pneumoniae* strain was isolated from the inner surface of an incubator. However, this strain was not stored for further research.

Screening and monitoring of extended spectrum beta-lactamase-producing and carbapenemase-producing *K. pneumoniae* in hospitals is crucial (21). However, the current molecular biological techniques, such as PCR and fluorescence quantitative PCR, have limitations and cannot be used extensively to screen for drug resistance genes. Genome sequencing maybe an effective method for screening resistant genes, although the technique entails high costs and is time-consuming, limiting it application. Homology analysis methods, such as ERIC-PCR, PFGE and MLST are complex and also time-consuming, such that timely detection of outbreaks is not feasible (22-24). Newer technologies need to be developed for monitoring the rapid outbreak of nosocomial infections.

Most previous reports on *K. pneumoniae* outbreaks involved single-site infections, diagnosed from respiratory specimens (25, 26), blood specimens (27-29), or urine specimens (30-32). This study showed that the pathogens were simultaneously isolated from respiratory specimens, blood specimens and umbilical vein catheter tips in the same patient. However, isolates of kpn1340, kpn1341, kpn1356, kpn1357, kpn1358, kpn1359 were not collected and could not be further analysed. Antibiotic susceptibility reports revealed that in all cases antibiotic susceptibility was consistent with strain ST37 or ST3006 and we suspect that these may belong to the same clones.

A number of limitations to our research should be noted. Specifically, we were not aware of a nosocomial infection outbreak; therefore, some strains of *K. pneumoniae* were not collected. It is important to enhance awareness of infection outbreak and strengthen bacterial preservation. Furthermore, we should strengthen communication and cooperation with infection control practitioners to screen and block nosocomial transmission at an earlier stage. Hospitals should implement different strategies to combat outbreaks, such as strict hand-washing by staff, frequent equipment change and extensive cleaning of paediatric wards.

## Conclusions

In summary, we reported two clones of *K. pneumoniae* ST37 and ST3006 outbreak in a neonatal intensive care unit. The clones were isolated from multi-location, sputum specimens, blood and umbilical vein catheter tips. Whole-genome sequencing would be a useful method for the detection of drug resistance genes and the analysis gene function. Active and effective infection control measures are indispensable for preventing and controlling nosocomial infection outbreaks.

## Acknowledgement

We thank the team of curators of the Institut Pasteur MLST and whole-genome MLST databases for curating the data and making them publicly available at http://bigsdb.pasteur.fr

## Ethics approval

The ethics approval number is K2018-01-001.

